# Efficient gastrointestinal colonization by *Campylobacter jejuni* requires components of the ChuABCD heme transport system

**DOI:** 10.1101/2025.03.18.643992

**Authors:** Vincent R. Randaisi, Madison L. Bunch, Will N. Beavers, Theresa Rogers, Ryan Mesler, Tiara D. Ashurst, Dallas R. Donohoe, Andrew J. Monteith, Jeremiah G. Johnson

**Affiliations:** The University of Tennessee Knoxville; The University of Iowa; Louisiana State University

## Abstract

Previous research demonstrated that *Campylobacter jejuni* encodes a heme utilization system that facilitates heme-dependent growth under iron-limiting conditions and that transcription of this system is induced during human infection. Despite these observations, it remained unknown whether the heme transport system is required for colonization and disease in a susceptible host. To address this, we created individual non-polar deletion mutants of each component of the heme transport system, as well as a total deletion of the inner membrane transporter, ChuBCD, and examined their ability to promote heme-dependent growth and iron uptake. From this work, we found that only the heme receptor, ChuA, was required for heme-dependent growth and iron acquisition, which supports earlier work of another group. Further, we examined whether intestinal colonization, immune activation, and pathology were altered during infection with these mutants. After establishing that elevated heme and *chuABCD* expression occurs during *C. jejuni* infection of IL-10^-/-^ mice, we found that heme transport mutants exhibited significantly reduced fecal shedding and colonization of the cecum and colon. In addition, we found that neutrophil and macrophage recruitment and intestinal pathology often remained intermediately elevated despite decreased bacterial loads. These results suggest that heme utilization promotes efficient colonization and full pathogenicity in *C. jejuni*, but that neither is completely abrogated in its absence.

## Introduction

Infection by *Campylobacter* species is a significant global health issue, causing an estimated 400 million cases of gastroenteritis each year, and resulting in substantial morbidity, mortality, and economic burden [1, 2]. Epidemiological studies have found that most individuals are infected with the species *C. jejuni,* and that they experience self-limiting moderate-to-severe gastroenteritis which is accompanied by inflammatory diarrhea and stomach pain [3-7]. In addition to acute gastrointestinal disease, campylobacteriosis is associated with long-term complications, including post-infectious irritable bowel syndrome, Guillain-Barré Syndrome, Miller-Fisher syndrome, and reactive arthritis [7-10]. Despite the impact of these infections on human health, little is known about how the host responds to *Campylobacter* spp. and how the bacterium counteracts or promotes host responses to facilitate colonization, disease, and/or dissemination.

Nutritional immunity is a defensive mechanism used by hosts to restrict access to vital nutrients to invading pathogens, including iron [11]. For example, iron in host tissues is often found in complexes with host molecules, including heme, lactoferrin, and transferrin, leading free iron to be essentially absent [12-14]. In response to infection, this restriction is further contributed to by the immune-mediated release of iron-uptake promoting molecules like hepcidin and leukocyte-derived iron sequestering proteins like calprotectin [15]. Relevant to gastrointestinal infection, several of these nutritional immunity factors are present in the gastrointestinal tract, where iron availability can be further reduced through complexing to various dietary components and microbiota-derived iron-scavenging molecules like siderophores [16, 17]. These processes combine to effectively limit growth and survival of pathogens in hosts and facilitate eventual clearance. In the case of campylobacteriosis, our group and others have demonstrated that nutritional immunity factors like calprotectin, Lcn-2, and S100A12 are significantly increased in the gastrointestinal lumen during human infection, which suggests the bacterium is restricted for iron [18, 19].

Pathogenic bacteria have evolved several mechanisms to counteract nutritional immunity, including the acquisition of iron directly from several host molecules, including heme, lactoferrin, and transferrin. In heme utilization, a significant amount of research has focused on elucidating these mechanisms in Gram-positive pathogens, including *Staphylococcus aureus,* or in Gram-negative pathogens, like *Pseudomonas aeruginosa* [20, 21]. For example, the Phu system in *P. aeruginosa* is used to acquire nutritional iron from heme and is comprised of, PhuA, an outer membrane receptor that binds hemoproteins like hemoglobin and myoglobin; the TonB-dependent energy transducer, PhuR, which interacts with PhuA to transfer heme across the outer membrane; PhuT, which is a periplasmic heme chaperone; and the ABC transporter PhuVW, which translocates heme across the inner membrane into the cytoplasm [22, 23]. In addition to the Phu system, *P. aeruginosa* uses the Has system for heme utilization which uses distinct gene products to perform heme uptake [24]. Similarly, *C. jejuni* encodes a heme utilization operon (*chuZABCD*) that was shown to facilitate growth in iron-restricted cultures when exogenous heme was added as an iron source (Figure 1) [25]. This study found that heme-dependent growth relied on the heme oxygenase, ChuZ, and the heme receptor, ChuA, but not the inner membrane ABC transporter, ChuBCD. In subsequent studies, *chuZABCD* was found to be upregulated in the feces of healthy volunteers infected during a human feeding study [26]. Because *chuZABCD* have been shown to be negatively regulated by the ferric-uptake regulator (Fur), this result suggests that the lower gastrointestinal tract of humans possesses low-levels of free iron and that the acquisition of heme-derived iron may be required for colonization and/or growth [27]. Importantly, it remains unknown whether the Chu heme translocation system (ChuABCD) facilitates *C. jejuni* infection of animals susceptible to disease and whether the inner membrane ABC transporter (ChuBCD) is as dispensable *in vivo* as it is *in vitro*.

**Fig. 1.**
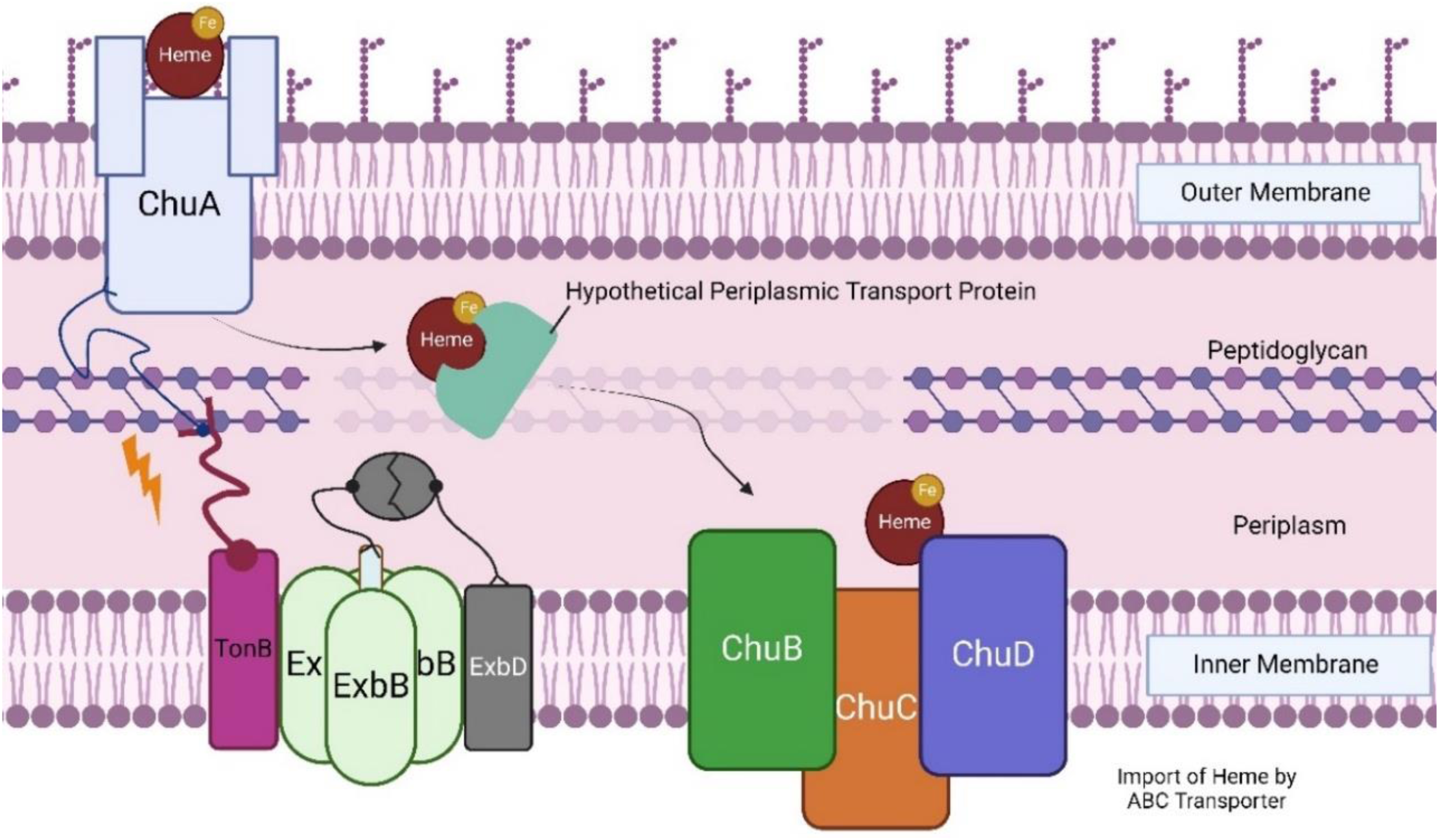
Proposed model of the *C. jejuni* Chu System. ChuA is a TonB-dependent outer membrane heme receptor with heme being transported into the periplasmic space via proton motive force. Once in the periplasm, heme may bind an unidentified periplasmic transport protein. Heme then associated with the inner membrane ChuBCD complex. ChuB is an inner membrane permease, ChuC is an ABC transporter, and ChuD is a periplasmic hemin-binding protein. In the cytoplasm, the heme oxygenase, ChuZ, then processes the porphyrin ring and releases iron for use in various metabolic processes (not shown).

To investigate whether heme utilization is required for gastrointestinal infection by *C. jejuni*, we constructed in-frame deletion mutants of the individual components of the transport system (ChuABCD) and examined their ability to acquire nutrient iron from heme under iron-restricted conditions. We found that only the heme receptor, ChuA, was required for heme utilization *in vitro* and that the presumed cognate ABC transporter, ChuBCD, was not required for this process. When we investigated the role of this system *in vivo*, we found that all determinants were required for efficient colonization and that, despite exhibiting reduced fecal and tissue burden, the mutants caused intermediate innate immune cell recruitment and gastrointestinal pathology.

## Results

### Only the predicted heme receptor, ChuA, is required for *in vitro* utilization of heme as an alternative iron source

In a previous study, insertion mutants of the ChuABCD transport system were examined for growth under iron-restricted conditions where heme was added as an alternative iron source and only growth of the ChuA mutant was significantly affected [25]. To confirm these earlier results, we constructed non-polar deletion mutants versus the insertion mutants used in the earlier study, and examined for growth under iron-restricted conditions where heme was added as an iron source. After 48 hours of growth, in the absence of heme and under iron-restricted conditions, our data showed that all strains exhibited a significant reduction in growth when compared to iron-replete conditions without added heme: wild type (7.29%), Δ*chuA* (0.628%), Δ*chuB* (7.89%), Δ*chuC* (12.23%), Δ*chuD* (7.36%), and Δ*chuBCD* (16.45 %) (Figure 2A). When heme was added to similar iron-restricted cultures and compared to iron-replete cultures, we found that wild-type, Δ*chuB*, Δ*chuC*, Δ*chuD*, and Δ*chuBCD* growth were restored with relative iron-replete values of 117.05%, 90.21%, 87.01%, 99.89%, and 102.58%, respectively (Figure 2A). The results from each mutant were not significantly different than the heme-dependent growth restoration that was observed for wild-type *C. jejuni.* In contrast, the addition of heme was unable to restore growth for Δ*chuA* with a relative iron-replete value of only 5.02%; this result was significantly reduced when compared to wild-type growth restoration and indicates that ChuA is a dominant determinant in the uptake and utilization of heme under iron-restricted conditions.

**Figure 2.**
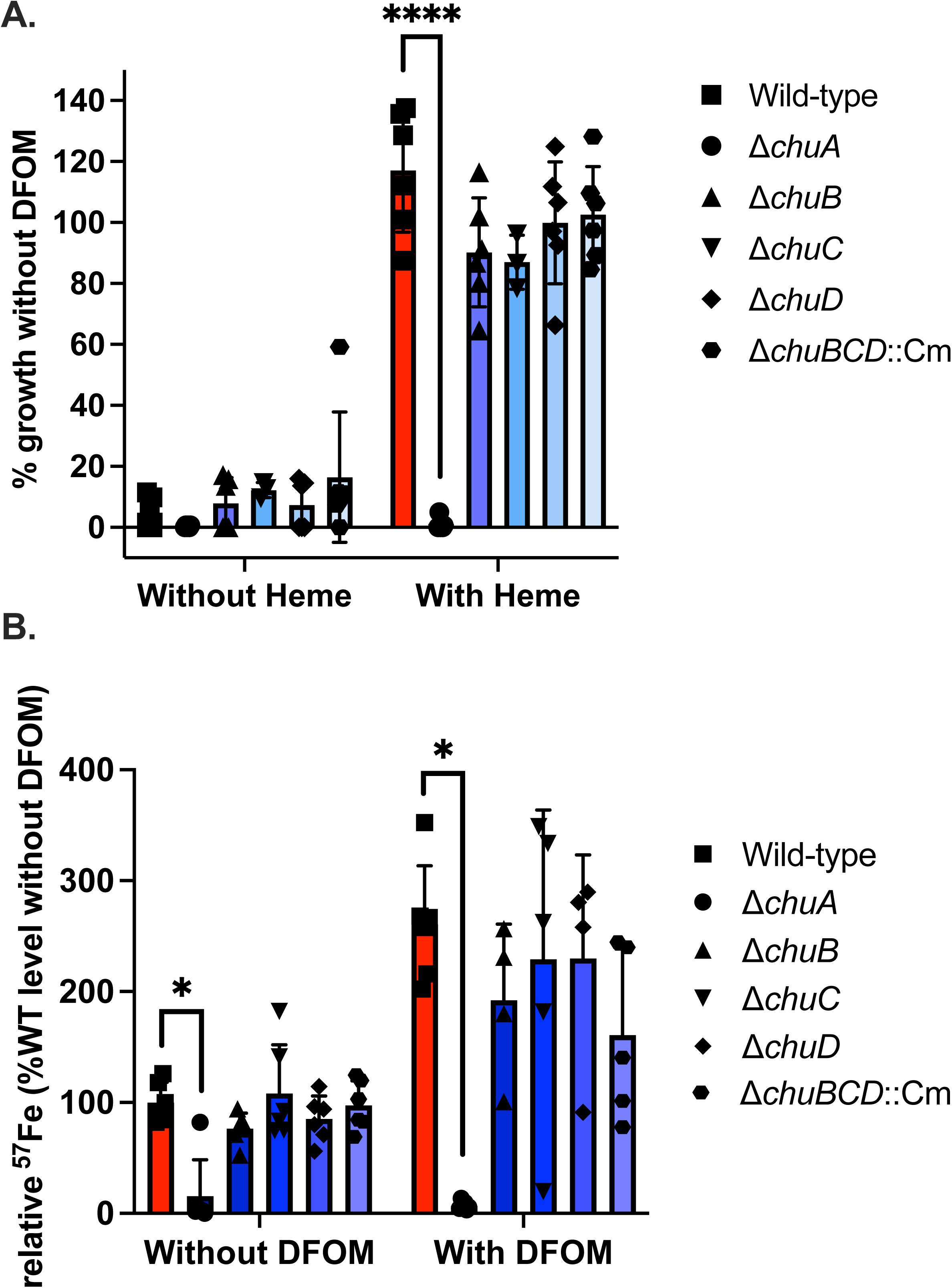
Growth and heme-dependent acquisition of iron in heme transport mutants under iron-restricted conditions. A) Wild-type, Δ*chuA*, Δ*chuB,* Δ*chuC,* Δ*chuD,* and Δ*chuBCD*::Cm were grown for 48 hours in MH broth with DFOM and with or without heme as an iron source. Presented as percent growth when compared to control cultures without DFOM added and are the result of two independent experiments with triplicate cultures. B) Wild-type, Δ*chuA*, Δ*chuB,* Δ*chuC,* Δ*chuD,* and Δ*chuBCD*::Cm were grown for 48h with or without DFOM in the presence of isotopically labeled heme (^57^Fe-heme) as an iron source. Presented as percent ^57^Fe when compared to control cultures without DFOM added and are the result of two independent experiments with triplicate cultures. \**p* < 0.05

To further verify the role of each determinant in heme uptake, we grew each deletion mutant under similar iron-restricted conditions but used ^57^Fe-heme to restore growth and measured the amount of cell-associated ^57^Fe for each strain. We first verified that DFOM and ^57^Fe-heme inhibited and restored growth, respectively, and compared the mutant strains to wild-type *C. jejuni* (Supp. Fig. 1). In MH media alone, all strains produced a similar number of CFUs to wild-type *C. jejuni* (Supp. Fig 1A). In MH media supplemented with DFOM, we successfully inhibited CFU production for each strain when compared to MH media alone, with average reductions between 9.4×10^-3^ to 9.0×10^-7^ though no strain was significantly impacted (Supp. Fig. 1B). In adjacent cultures, we added ^57^Fe-heme at the concentration used above to restore growth in the presence of DFOM and found that only Δ*chuA* exhibited a significant reduction in CFUs with a mean decrease of 9.4×10^-6^ (Supp. Fig. 1C). Using ICP-MS, we quantified the amount of cell-associated ^57^Fe for each strain under these conditions and found that only Δ*chuA* exhibited a significant decrease when compared to wild-type *C. jejuni* with an average concentration of 6.9% versus 260.6% (Fig. 2B). This result not only supports our earlier conclusion that ChuA is involved in acquiring iron from heme under iron-limited conditions, but because we observed a significant increase in ^57^Fe under iron restriction in wild-type *C. jejuni,* the Chu system is likely induced during iron restriction. Since this reduction could be due to inhibited growth of Δ*chuA*, we added ^57^Fe-heme to MH media alone and examined growth and the accompanying cell-associated ^57^Fe levels. From this analysis, none of the strains were found to exhibit a significant difference in CFUs when compared to wild-type *C. jejuni* (Supp. Fig. 1D). When we quantified ^57^Fe levels using ICP-MS, we found that only Δ*chuA* exhibited a significant reduction when compared to wild-type *C. jejuni,* with average concentrations of 15.69% and 100%, respectively. This supports the above result that only ChuA is involved in the *in vitro* acquisition of iron from heme and that the Chu system is likely expressed at a basal level under iron-replete conditions since strains containing ChuA still secured significant amounts of ^57^Fe from heme.

### Heme and *chu* expression increase during *C. jejuni* infection

Campylobacteriosis in humans often leads to inflammatory diarrhea accompanied by blood in the stool. To quantify the effect of infection on fecal heme accumulation in humans, we leveraged surveillance stools collected by our group from healthy, uninfected volunteers, as well as clinical samples from patients known to be infected with *Campylobacter* and no other gastrointestinal pathogen. We used HPLC to measure fecal heme and found that feces from uninfected volunteers contained 2.90 pmol/mg and the feces from infected patients possessed significantly more heme at 753.83 pmol/mg, a 260-fold increase (Fig. 3A). This observation suggests that heme is present at elevated levels during *Campylobacter* infection and could serve as a nutrient source, which is further supported by a previous study where *chuZABCD* expression significantly increased during *C. jejuni* infection of healthy human volunteers [24]. To establish whether similar responses are observed in the mouse model of campylobacteriosis, we collected feces from mice infected with wild-type *C. jejuni* and quantified heme by HPLC. For heme, we observed mean concentrations of 1.82 pmol/mg of feces for uninfected mice and 3.64 pmol/mg of feces for animals infected with wild-type *C. jejuni* (Fig. 3B). This two-fold increase in fecal heme represented a significant accumulation when compared to uninfected animals. To examine whether the bacterium expresses the Chu system to acquire heme during murine infection, we extracted RNA from *in vitro* grown iron-replete and iron-restricted media, cecal tissue, colon tissue, cecal contents (chyme), and feces, and quantified *chuA* transcript abundance relative to iron-replete, *in vitro* grown cells. From this analysis, we found that *chuA* transcript abundance significantly increased during iron-restriction *in vitro,* with a mean increase of 28.71-fold. Similarly, c*huA* abundance was significantly increased in all mouse tissue with fold-changes of 150.48 for cecum, 823.92 for colon, 163.63 for chyme, and 384.77 for feces when compared to iron-replete, *in vitro* grown cells (Fig. 3C). When comparing these values, the 5.48-fold increase observed in the colon when compared to the cecum, as well as the 2.35-fold increase in the feces when comparing to the chyme, were significant. These changes suggest that during infection, *C. jejuni* increases expression of the Chu system to gain access to the heme that is released from the host, particularly in the colon and feces.

**Figure 3.**
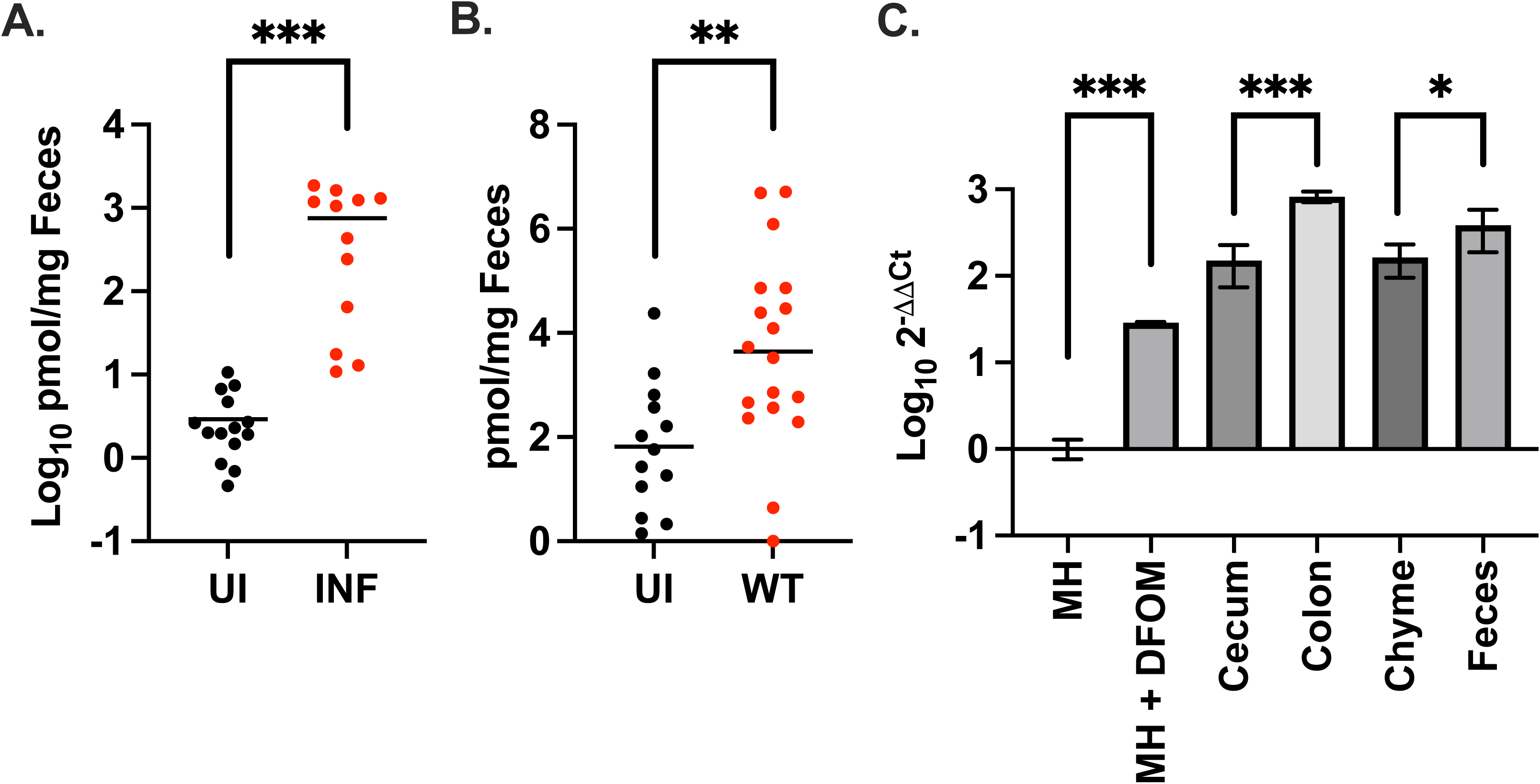
Fecal heme and expression of the Chu heme transport system during infection. A) Levels of heme b were measured by mass spectrometry in the feces of uninfected (UI) volunteers or in clinical samples from patients infected with only *Campylobacter* (INF). B) Levels of heme b were measured by mass spectrometry in the feces of uninfected (UI) IL-10^-/-^ mice or in the feces of IL-10^-/-^ mice infected with wild-type *C. jejuni*. C) RT-qPCR analysis of *chuA* expression in wild-type *C. jejuni* grown in iron-replete MH (MH) or iron-restricted MH (MH+DFOM), or in the cecum, colon, chyme, or feces, during infection of IL-10^-/-^ mice. Expression relative to the *C. jejuni* control gene, *rpoA.* \**p* < 0.05; \*\**p* < 0.01; \*\*\**p* < 0.001

### All Chu determinants are required for efficient colonization of mice susceptible to *C. jejuni* infection

To determine how each component of the heme transport system affects *C. jejuni* infection, each of the deletion mutants was used to orally inoculate IL-10^-/-^ mice that are susceptible to colonization and *C. jejuni-*induced gastrointestinal inflammation. Feces were collected every two days for up to 10 days when the animals were euthanized and tissues harvested. Wild-type *C. jejuni* consistently colonized mice that was characterized by a steady increase in fecal CFUs that leveled-out at six days post-infection and remained high throughout the 10-day experiment. In contrast, fecal homogenates from uninfected mice did not yield detectable CFUs throughout the 10-day study. Compared to wild-type, Δ*chuA* was significantly reduced throughout the study at days 2, 6, 8, and 10 post-infection while Δ*chuB* was only significantly reduced at days 6 and 8 post-infection. Like Δ*chuA,* the Δ*chuC* and Δ*chuD* mutants were significantly reduced at all timepoints with an apparent inability to reach the levels of colonization observed for wild-type. Lastly, the Δ*chuBCD* mutant was only significantly reduced at days 2 and 10 post-infection (Supp. Fig. 2).

When examining the cecal tissue at 10 days post-infection, wild-type *C. jejuni* exhibited the highest amount of cecal colonization, with a mean of 1.93 x 10^7^ CFU/g of tissue. By comparison, each mutant strain except Δ*chuBCD* exhibited significantly reduced cecal colonization with means of 5.86 x 10^6^ CFU/g for Δ*chuA,* 3.72 x 10^5^ CFU/g for Δ*chuB,* 2.88 x 10^5^ CFU/g for Δ*chuC,* 1.92 x 10^6^ CFU/g for Δ*chuD*, and 3.25 x 10^6^ CFU/g for Δ*chuBCD* (Fig. 4A). In addition, CFUs were quantified for homogenized colons with wild-type, Δ*chuA,* Δ*chuB,* Δ*chuC,* Δ*chuD,* and Δ*chuBCD* exhibiting mean colonization loads of 9.72 x 10⁶ CFU/g, 9.06 x 10^4^ CFU/g, 3.91 x 10⁶ CFU/g, 7.14 x 10^5^ CFU/g, 1.71 x 10^5^ CFU/g, and 2.36 x 10^5^ CFU/g, respectively (Fig. 4B). When compared to mice infected with wild-type *C. jejuni*, these results indicate that each mutant is significantly reduced in its ability to colonize the colon. Lastly, the bacterial loads in fecal samples corresponding to this timepoint were quantified (above). Wild-type *C. jejuni* exhibited the highest average colonization with a mean of 9.47 x 10^7^ CFU/g while the mutant strains colonized at 2.02 x 10^5^ CFU/g for Δ*chuA*, 5.63 x 10^7^ CFU/g for Δ*chuB*, 2.54 x 10^7^ CFU/g for Δ*chuC*, 7.00 x 10^6^ CFU/g for Δ*chuD*, and 7.15 x 10^5^ for Δ*chuBCD* (Fig. 4C). These results suggest that each mutant, except Δ*chuC,* are significantly decreased for viable CFUs in the feces when compared to wild-type *C. jejuni.* Taken together, it appears that despite some mutations not being affected depending on the region of the gastrointestinal tract examined, the Chu determinants are consistently required for full colonization of the gastrointestinal tract.

**Figure 4.**
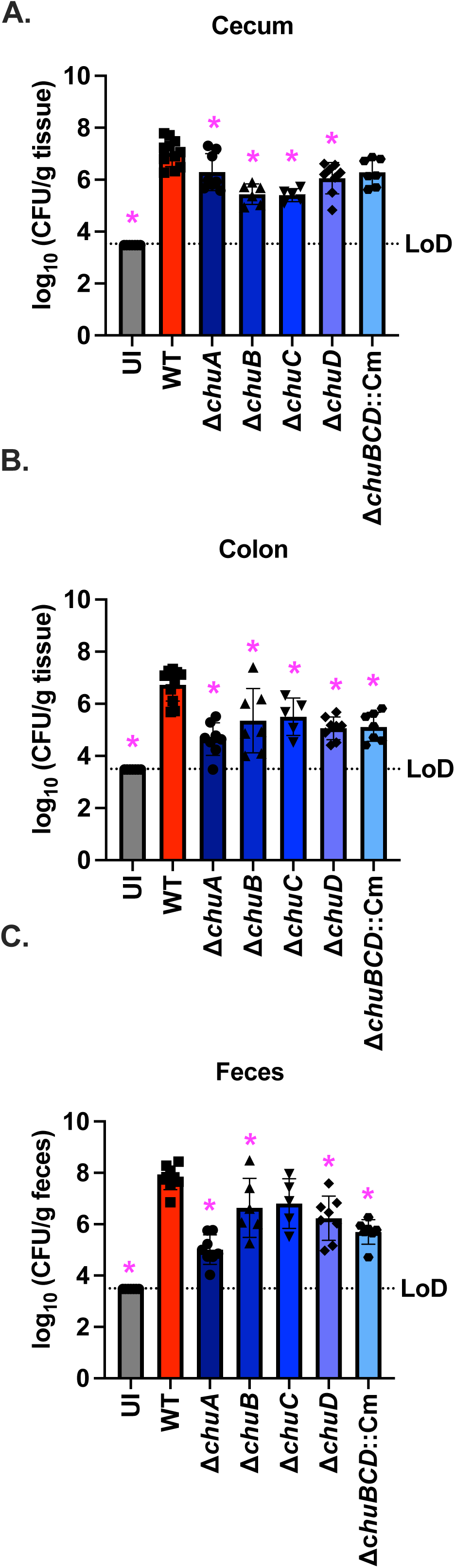
Colonization of mice by mutants of the Chu heme transport system. *C. jejuni* burden was measured at ten days post-infection by calculating the Log_10_ transformed CFU/gram of wild-type and each mutant in the cecum (A), colon (B), and feces (C). \**p* < 0.05

### Chu system mutants cause intermediate immune cell recruitment and significant intestinal pathology despite reduced colonization

Macrophage and neutrophil recruitment to the cecum and colon of infected mice was quantified by immunofluorescence microscopy using specific markers for each cell type. From this analysis, the cecal tissue generally contained more macrophages and neutrophils than the colon even in an uninfected animal. When comparing immune cell recruitment between infected animals, all exhibited significant increases in macrophages and neutrophils in both the cecum and colon when compared to an uninfected animal except for macrophage recruitment to the cecum of Δ*chuD* and Δ*chuBCD* infected mice (Fig 5A-D). In contrast, when immune cell recruitment was compared to a wild-type-infect animal, Δ*chuA* was significantly reduced for neutrophils in the cecum and colon (Fig. 5AB), and macrophages in the colon (Fig. 5D). In addition, Δ*chuD* and Δ*chuBCD* were decreased for neutrophils and macrophages in the cecum, and macrophages in the colon (Fig. 5ACD). While macrophage recruitment to the cecum during infection with Δ*chuA* was not significantly reduced when compared to a wild-type infected mouse, there is a trend toward reduced macrophage recruitment to the colon (p=0.058). This result suggests that immune cell recruitment is consistently reduced during infection with Δ*chuA,* Δ*chuD,* and Δ*chuBCD*.

**Figure 5.**
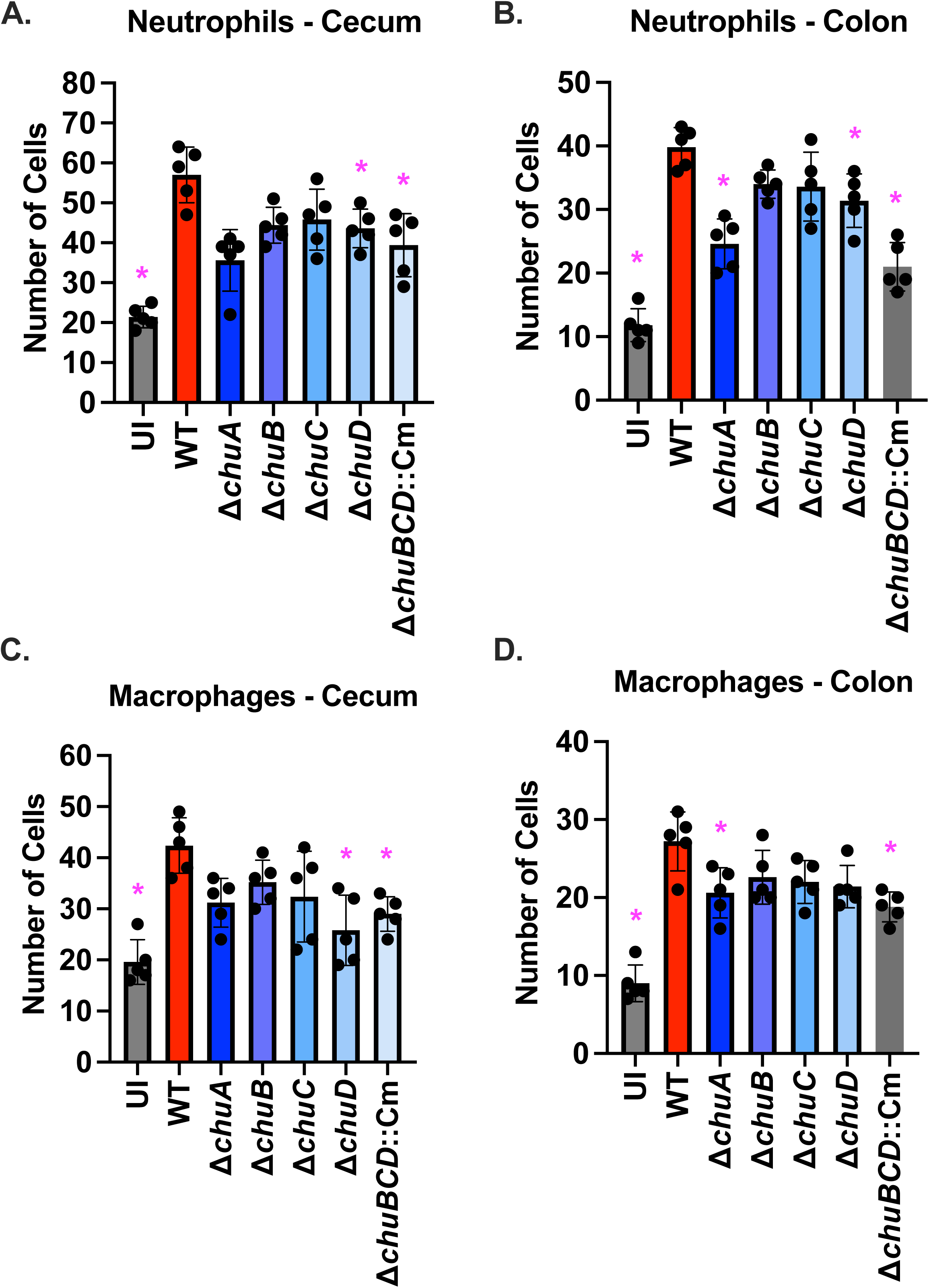
Recruitment of innate immune cells to gastrointestinal tissue during infection with mutants of the Chu heme transport system. Immunofluorescence microscopy of neutrophils and macrophages were counted in fixed and embedded tissue from an uninfected IL-10^-/-^ mouse or those infected with either wild-type *C. jejuni* or each Chu heme transport system mutant. The number of neutrophils in the cecum (A) or colon (B) and macrophages in the cecum (C) or colon (D) were statistically compared to the number observed during wild-type infection. \**p* < 0.05

Because of decreased innate immune cell recruitment in the cecum and colon, we evaluated the pathology of intestinal tissues in mutant infected mice. When examining for edema in the cecum and colon, we found that uninfected animals exhibited significantly reduced swelling in both tissues when compared to wild-type infected animals (Fig. 6A). All other mutants and tissues were indistinguishable from wild-type infected animals. Similarly, uninfected animals had significantly less hyperplasia of the cecum and colon when compared to wild-type infected animals, but only the Δ*chuC* strain was found to have significantly less hyperplasia in the cecum (Fig. 6B). The most consistent difference observed among the mutants was in goblet cell loss, where uninfected animals and all mutant infected cohorts exhibited significantly higher numbers of goblet cells in the cecum and colon when compared to wild-type infected animals (Fig. 6C). When comparing uninfected animals and mutant infected groups, we similarly found that all mutant cohorts had significantly higher numbers of goblet cells. This indicates that mutant infected animals exhibit an intermediate response in terms of goblet cell numbers where they possess more than wild-type infected animals, but less than uninfected mice.

**Figure 6.**
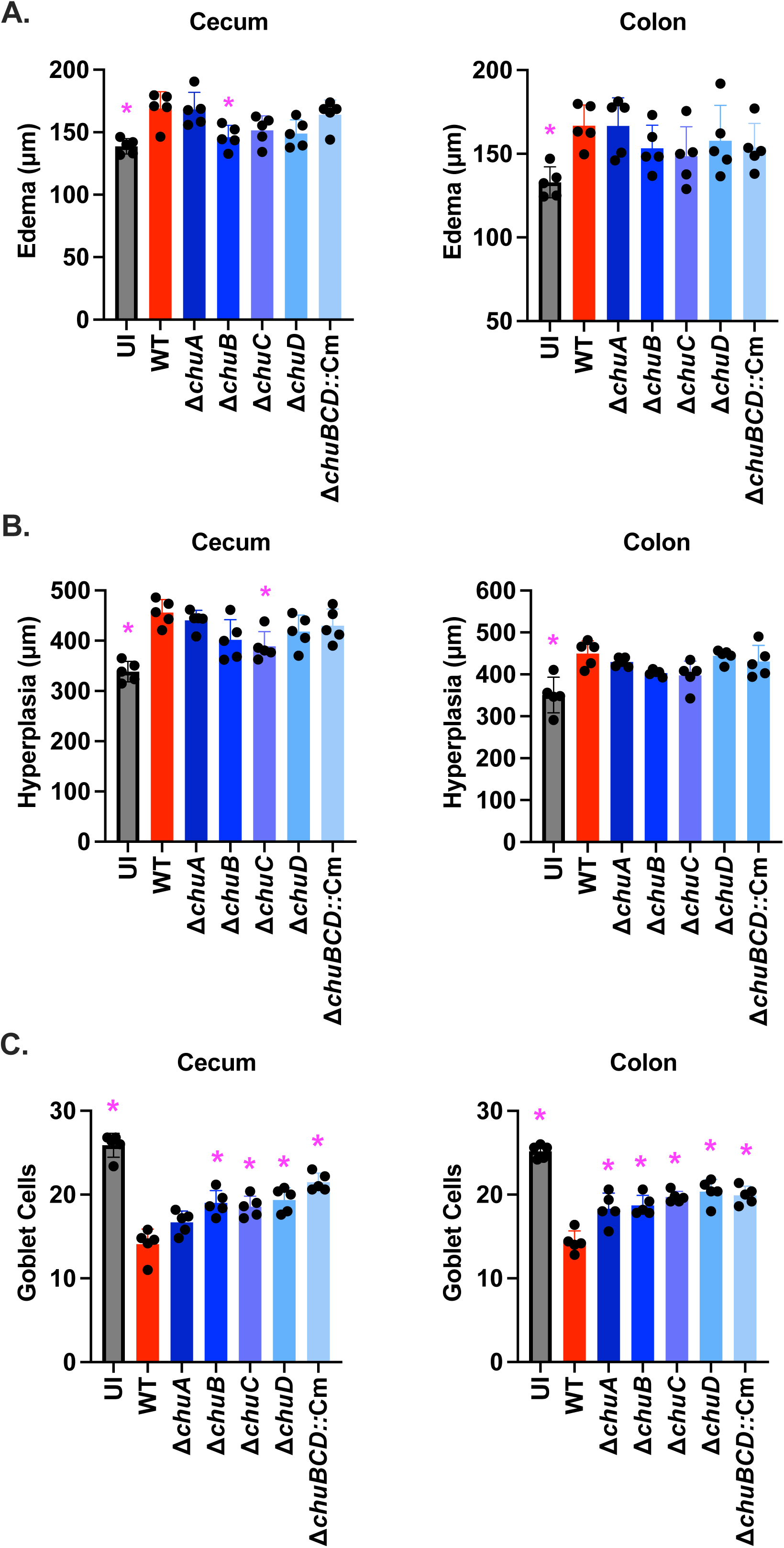
Histopathology of gastrointestinal tissues during infection with mutants of the Chu heme transport system. Multiple characteristics were measured in a blinded manner on H&E-stained, fixed and embedded tissue from uninfected IL-10^-/-^ mice or those infected with either wild-type *C. jejuni* or each Chu heme transport system mutant. Edema (A), hyperplasia (B), and goblet cell loss (C) were measured in the cecum and colon for each infection and statistically compared to the numbers recorded during wild-type infection. \**p* < 0.05

## Discussion

Supporting earlier work, we found that only ChuA was needed for growth when heme was added as an alternative iron source under iron-restricted conditions, since individual mutants and a triple mutant of the predicted inner membrane transporter were able to grow at levels indistinguishable from wild-type under those conditions. Going a step further, we supplemented similarly restricted cultures with isotopically labeled heme and quantified cell-associated levels of that isotope and found that ChuA was the only determinant required for acquisition of iron from heme. Further, this experiment demonstrated that acquisition of heme-derived iron increased under iron restriction, which supports gene expression studies showing *chuZABCD* transcript abundance increases under low iron availability due to control by the ferric uptake regulator (Fur) [27]. We also found that ChuA-dependent acquisition of heme-derived iron could be detected under iron-replete condtions, which suggests there is some level of constitutive Chu expression. When we considered why ChuA is the only determinant required for heme utilization, we initially hypothesized there is redundancy in the genome for ChuBCD. This hypothesis was supported by the observation that there is no apparent homolog of ChuA in the *C. jejuni* 81-176 genome and that CeuB, the ABC-type enterochelin permease, exhibits some homology to ChuB: 30% (94/310) amino acid identity and 50% (157/319) positive residues. In addition, the ABC-type enterochelin ATPase, CeuD, exhibits some homology to ChuC: 30% (72/242) amino acid identity and 58% (142/242) positive residues. Beyond CeuD, an additional 22 genes encode potential ChuC homologs with 92-73% coverage and 36.97-24.31% amino acid identity. An observation that did not support the hypothesis of functional redundancy is that no apparent homolog of ChuD exists in the *C. jejuni* 81-176 genome, though the functional annotation of ChuD and CeuE are the same: ABC transporter periplasmic binding component. Additional work will need to be conducted to establish whether components of the enterochelin transport system can functionally substitute for Chu determinants.

To examine the contribution of each determinant to animal infection, we first infected IL-10^-/-^ mice with wild-type *C. jejuni* and quantified the abundance of heme in the feces. From this experiment, we found that wild-type infected animals exhibited elevated amounts of heme in their feces. Interestingly, the release of heme was far less than what was observed in human volunteers and patients, where uninfected individuals had fecal heme that surpassed even that of infected mice. It is unclear why the amount of heme between the species is drastically different, but it is likely the result of both dietary heme consumption and propensity to develop blood in the stool during infection. In terms of dietary heme, uninfected volunteer fecal heme was markedly higher than what was observed in any mouse cohort, perhaps due to meat consumption by volunteers in contrast to the predominantly plant-based diet of laboratory mice. In the case of susceptibility to bleeding, it was apparent from this analysis that humans develop more pronounced hematochezia than even IL-10^-/-^ mice during campylobacteriosis. Finding the mechanism behind this difference may lead to the mechanism behind why campylobacteriosis is more severe in humans than other hosts.

We also found that *C. jejuni* significantly increases Chu system expression in the colon and feces when compared to the cecum and cecal contents, though all were elevated relative to *in vitro* grown bacteria, including cultures treated with DFOM. This result suggests: i) the animal gastrointestinal tract is a potent inducer of the heme utilization sytem, ii) the colon and feces may be more iron limited than the cecum and chyme since low iron induces Chu system expression, and iii) *C. jejuni* likely takes advantage of heme as an alternative iron source in all regions of the gastrointestinal tract during infection. When we examined for how mutation of each Chu determinant impacted colonization of IL-10^-/-^ mice, we found that almost all were significantly reduced in the cecum, colon, and feces at 10 days post-infection relative to wild-type infected mice. This was interesting to us because it suggests that heme acquisition *in vivo* partially relies on ChuBCD, which was not the case in our *in vitro* experiments. Why ChuBCD appear to be uniquely required for *in vivo* heme acquisition and colonization is unknown, but it may support the hypothesis that enterochelin transporters are co-opted for heme uptake during *in vitro* growth. *C. jejuni* does not synthesize its own enterochelin and likely relies on the production of the molecule by commensal bacteria during animal infection [28]. During *in vitro* growth where enterochelin transport is induced by iron restriction, these components become more abundant in the absence of their cognate ligand, due to monoculture, and may then be used for off-target applications like heme uptake. In contrast, during gastrointestinal infection, siderophores like enterochelin may become available and be transported by this dedicated system, thus creating competition for its use and greater dependence on the ChuBCD components for heme uptake. This bottleneck could then contribute to less efficient iron acquisition during infection with the mutants and lower colonization potential.

Further, we did not observe a drastic improvement in the intestinal pathology of mutant infected mice despite the bacterial burden being significantly lower with both edema and hyperplasia remaining elevated. The only consistent improvement in intestinal health was the intermediate increase of goblet cells in mutant infected mice when compared to wild-type-infected animals. A reduction in goblet cells appears to be generally observed in IL-10^-/-^ mice due to the role of the cytokine in goblet cell development and function, which can be exacerbated during chemical- or infection-induced colitis in both wild-type or IL-10^-/-^ mice. For example, goblet cell loss was observed during infection of wild-type mice with *C. rodentium* and although some of the goblet cells were directly infected with the bacterium, localization did not correlate with loss, indicating goblet cell loss is due to host-mediated inflammation [29]. This is supported by our observation that we observed both intermediate innate immune cell recruitment and loss of goblet cells in response to Chu mutant infection. This result suggests to us that goblet cell loss may be a more sensitive indicator of gastrointestinal inflammation than edema and hyperplasia. In addition, goblet cell loss may also be partly responsible for elevated edema and hyperplasia, despite reduced colonization, since the associated reduction in the mucus barrier can negatively impact gastrointestinal health.

It is intriguing to envision how developing strategies that combat heme utilization in the gastrointestinal tract may counteract *C. jejuni* infection. For example, if we could discover how *C. jejuni* causes intestinal bleeding to liberate this nutrient source, it is conceivable that we can increase intestinal integrity and health while reducing colonization at the same time by inhibiting that process. In addition, we could use more bacteria-centric approaches, as done with other pathogens, and identify molecules that can specifically inhibit heme utilization components, like ChuZABCD. This is an interesting proposition since, as mentioned above, *C. jejuni* does not synthesize its own siderophores and may naturally find itself in a competitive space for iron during infection. Because of this, developing anti-heme utilization strategies may be particularly useful in combating *C. jejuni* infection when compared to their use against pathogens that possess a greater array of iron acquisition systems, like *E. coli* and *Salmonella* [30, 31]. Before addressing these possibilities, it will be necessary in the future that we address the questions above relating to the use of enterochelin systems to transport heme, whether competition with enterochelin-producing commensals contributes to *C. jejuni’*s dependence on heme during infection, and whether direct inhibition of the heme utilization components in *C. jejuni* can reduce colonization.

## Material and Methods

### Bacterial Culture and Growth Conditions

The *Campylobacter jejuni* 81-176 strain DRH212 (wild-type) was stored at -80°C in Mueller-Hinton broth supplemented with 20% glycerol. All *C. jejuni* strains were routinely cultured on *Campylobacter-*selective (CS) medium containing 10% sheep’s blood, cefoperazone (40 µg/mL), cycloheximide (100 µg/mL), trimethoprim (10 µg/mL), and vancomycin (100 µg/mL). Cultures were incubated at 37°C under microaerobic conditions (85% N_2_, 10% CO_2_, and 5% O_2_) for 24-72 hours. *E. coli* strains used in this study were stored at -80°C in low-salt LB broth supplemented with 20% glycerol. These strains were regularly grown aerobically at 37°C in low-salt LB broth or on low-salt LB agar plates. When needed, ampicillin was added at a final concentration of 100 µg/mL.

### Mutant Generation

To create *chu* mutants, approximately 500 basepairs up and downstream of each gene were amplified by PCR and stitched together using splicing by overlap extension PCR (SOE-PCR). Each stitched fragment was cloned into pJET1.2 vector using the manufacturer’s protocol (ThermoFisher Scientific, Cat. # K1231). Transformants were selected on low-salt LB agar with ampicillin and grown at 37°C overnight. Colonies were screened for the insert by PCR and sequenced. Correct plasmids were digested with BamHI and a similarly cut *rpsL*::*cat* cassette was ligated between the upstream and downstream regions of each gene and transformants were selected on LB with chloramphenicol (15 µg/mL). Colony PCR was performed to confirm insertion and plasmids were sequenced. Each plasmid was electroporated into wild-type *C. jejuni* and transformants were selected on media containing chloramphenicol (15 µg/mL). Deletion of the target gene and insertion of *rpsL*::*cat* was confirmed by PCR. These insertion-deletion mutants were used to create clean deletion mutants by electroporating in the original pJET-SOE-PCR fragment without the *rpsL*::*cat*. Chloramphenicol-sensitive, streptomycin resistant colonies were screened for target gene deletion by PCR and in-frame deletion was confirmed by whole-genome sequencing. Despite repeated attempts, we were unable to conctruct an in-frame deletion mutant of *chuBCD* and proceeded with the intermediate strain containing the *rpsL*::*cat* cassette (Δ*chuBCD*::Cm). PCR and whole-genome sequencing confirmed the insertion of *rpsL*::*cat* and deletion of the *chuBCD* genes in this mutant.

### Heme Growth Assay

Wild-type and Δ*chu* mutants were inoculated on CS agar for 24h then passed onto another CS agar plate and grown for 24h. Using a 96-well plate format and analyzing samples in triplicate. Cells were harvested into MH broth to an OD A600 of 0.05. One hundred microliters of suspension was added to either i) an equal volume of MH broth, ii) an equal volume of MH broth containing 200 μM of hemin, iii) an equal volume of MH broth containing 80 μM of deferoxamine mesylate (Sigma-Aldrich, Cat. # 252750), or iv) an equal volume of MH broth containing 200 μM of hemin and 80 μM of deferoxamine mesylate (DFOM). Due to the addition of an equal volume of culture, the final concentration of each reagent was halved. An uninoculated control was also made for each experimental condition. Strains were incubated in microaerobic conditions overnight at 37°C and bacterial growth was quantified at OD A600. Each strain was assayed in triplicate at least three times to ensure reproducibility and statistical analysis performed using a one-way ANOVA.

### 57Fe Growth Experiments

*C. jejuni* strains were grown on CS media as above and used to start a 5 mL overnight culture in MH broth. Cells were used to inoculate 5 mL of MH broth in optically clear tubes at an OD A600 of 0.025 that contained either i) MH alone, ii) MH with 10µM DFOM, iii) MH with 100 µM ^57^Fe-heme (Fisher Specialty Chemicals, Cat. # P40080), or iv) MH with 100µM ^57^Fe-heme and 10µM DFOM. These cultures were incubated microaerobically at 37°C until mid-log phase (approximately 28h incubation). Cells were pelleted by centrifugation at 4000 rpm for 8 minutes, and the supernatant was discarded. The bacterial pellet was resuspended in water and centrifuged twice to ensure thorough washing. The final pellet was weighed, resuspended in 500 µL of water, and a sample was taken for serial dilution. The remaining suspension was added to a 15 mL conical tube containing 1 mL of 50% nitric acid and incubated overnight at 50°C on a heat block. The following day, 9 mL of water was added to the tube, and the sample was sent to Vanderbilt University for ICP-MS analysis. For serial dilutions, each was spread onto CS agar plates, incubated microaerobically at 37°C for 48-72 hours, and colonies counted. Statistical analysis was performed using a one-way ANOVA.

### Mouse Infection

All animal protocols were approved by the Institutional Animal Care and Use Committee at the University of Tennessee–Knoxville (UTK IACUC protocol #2885). Wild-type and the *chu* mutants were grown on CS medium and Gram stained to ensure culture purity before inoculation. Bacterial suspensions were made in sterile 1x PBS at approximately 10^10^ CFU/mL and used to oral gavage 8-to-10-week-old female IL-10^-/-^ C57BL/6 mice with 10^9^ CFUs of wild-type or the Δ*chu* mutants. In addition, female mice were mock infected with sterile 1x PBS. Fecal pellets were collected every two days for 10 days, weighed, diluted 1:10 in PBS, and homogenized. Fecal homogenates were serially diluted in 1x PBS and plated on CS media. Plates were incubated at 37°C under microaerobic conditions for 72 hours before CFUs were counted. Ten days after infection, mice were euthanized, and ceca and colons harvested and washed in sterile 1x PBS. A portion of each cecum and colon was placed in 10% buffered formalin for histology. The other portion was weighed and diluted 1:100 in PBS, and homogenized. Samples were serially diluted in sterile 1x PBS, plated on CS medium, and incubated for 72 hours before CFUs were counted. All measurements were recorded, and statistical analysis was performed using a one-way ANOVA.

### Heme extraction

Fecal pellets were collected at 10 days post-infection. Human feces or clinical specimens were collected in accordance with UTK IRB-17-03795-XM and IRB-23-07446-XP. All samples were weighed and resuspended in PBS. Heme was extracted three times with a volume of 1 N acetic acid in ethyl acetate equal to the PBS and homogenized through extensive vortexing. Organic/aqueous solvent phases were separated by centrifugation at 4000 rpm at 4°C for 10 min. Ethyl acetate layers were combined by transferring to clean glass tubes and dried using nitrogen gas. Each sample was subsequently dissolved in 100 µL of methanol for LC-MS/MS analysis. A heme standard was prepared in methanol at 1, 0.1, and 0.01 µM to serve as an external calibration control for quantification.

### LC-MS/MS analysis

The samples were analyzed at the LSU School of Veterinary Medicine Mass Spectrometry Resource Center on a Shimadzu 8060NX triple quadrupole mass spectrometer interfaced with a Shimadzu Nexera XS 40 series UHPLC and Shimadzu CTO-40S column oven. 5 µL of each sample, blank, or standard mixture was analyzed in positive ionization mode. Chromatographic separation was achieved using a Shimadzu Nexcol C 18 (50 mm length, 2.1 mm internal diameter, 1.8 µm particle size) column with a Phenomenex Security Guard C 18 (2.1 mm internal diameter) guard column. Mobile phase A was 0.1 % formic acid in water and mobile phase B was 0.1 % formic acid in acetonitrile. The column oven was set to 25 °C and the flow rate was set to 0.4 mL/min for the entire analysis. The starting condition was 40 % B, which was held for 1 min. The B mobile phase was then ramped to 98 % over the next 14 min. The column was washed at 98 % B for 5 min, then equilibrated to 40 % B for 5 min. All instrument voltages were determined and optimized empirically prior to the analysis. The m/z transitions were 616.3 to 557.3 for heme. An external calibration line of heme was used to quantify each sample before normalizing to fecal pellet weight. Statistical analysis was performed using a Mann-Whitney U-test.

### Immunofluorescence microscopy and enumeration of innate immune cells

Four-micron sections of each tissue were deparaffinized and antigens retrieved by microwaving. Tissues were washed with PBS and blocked for 30 minutes in normal rabbit serum (NRS) and normal goat serum (NGS). Samples were washed with PBS and stained at 4°C overnight with either Alexa 555 anti-CD11b antibody (1:100) or Alexa 647 anti-F4/80 antibody (1:50). Tissues were washed three times with PBS and mounted with VECTASHIELD antifade media containing DAPI (Vector Laboratories, Cat. # H-1200-10). The entire tissue section was imaged at the Central Microscopy Research Facility at the University of Iowa Carver College of Medicine. Neutrophils (Cd11b^+^) and macrophages (F4/80^+^) were counted in five fields-of-view for each animal and compared to the number recorded for a wild-type infected animal. Statistical analysis was performed using a one-way ANOVA.

### Histology and Slide Preparation

Pathology scoring for intestinal tissue was performed as previously described [32, 33]. Briefly, approximately one centimeter of the cecum and distal colon was removed, and the lumen was washed with sterile 1x PBS. The tissue was placed in 10% buffered formalin and fixed for 4 hours at room temperature. Fixed tissue was embedded in paraffin, and 4 μm sections were stained with hematoxylin and eosin (H&E). Stained slides were visualized using bright-field microscopy, and representative images were taken for pathology scoring. All imaging and scoring were performed blinded, and images were analyzed for edema, hyperplasia, and loss of goblet cells using ImageJ. Edema was assessed by measuring the interstitial space of each tissue. Goblet cells were identified, and the total count was recorded for each tissue section. Hyperplasia was measured by measuring the height of the intestinal epithelium from the base of the IEC using line measurement. All measurements were recorded, and statistical analysis was performed using a one-way ANOVA.

## Acknowledgements

This research was supported by NIH funds: K22AI153677 to W.N.B., R35GM154838 to A.J.M., and R01AI166535 to J.G.J. Research was also supported by USDA funds to D.R.D. and J.G.J.: NIFA-AFRI-2019-67017-29261. The University of Tennessee provided funds to A.J.M., including start-up and Human Health and Wellness Initiative funds. The acquisition of the Zeiss LSM 980 Airyscan2 Laser Scanning Confocal microscope that was used for fluorescent imaging of tissues was made possible by a generous grant from the Roy J. Carver Charitable Trust. Additional funding was provided by the University’s Office of the Vice President for Research. The authors would like to thank Dr. Wade Calcutt at the Vanderbilt University Mass Spectrometry Research Center and Jianqiang Shao at the University of Iowa Central Microscopy Research Facility for their assistance with the various analyses.

## Figure Legends

**Supplemental Figure 1. Colony forming units from ^57^Fe-heme utilization experiment.** CFUs were enumerated for each mutant under four conditions from three cultures grown on two independent occasions, including MH alone (A), MH with DFOM (B), MH with DFOM supplemented with ^57^Fe-heme (C), or MH with ^57^Fe-heme alone (D). * *p* < 0.05

**Supplemental Figure 2. Colony forming units from the feces of infected mice over ten days.** The number of CFUs present in fecal homogenates collected every other day from at least five individual mice infected with each heme transport mutant over the course of a ten-day infection.

